# Pre-existing intratumoral CD8 T cells substantially contribute to control tumors following therapeutic anti-CD40 and polyI:C based vaccination

**DOI:** 10.1101/2020.08.31.275420

**Authors:** Aaron D. Stevens, Timothy N.J. Bullock

## Abstract

**Background:** Dendritic cells are potently activated by the synergistic action of CD40 stimulation in conjunction with signaling through toll like receptors, subsequently activating antigen specific T cells. Cancer vaccines targeting the activation of dendritic cells in this manner show promise in murine models and are being developed for human cancer patients. While vaccine efficacy has been established, further investigation is needed to understand the mechanism of tumor control and how vaccination alters tumor infiltrating immune cells.

**Methods:** Mice bearing established murine melanoma tumors were vaccinated with agonist anti-CD40, polyI:C, and tumor antigen. Intratumoral T cell numbers, differentiation state, proliferation, and survival were assessed by flow cytometry. T cell effector function was measured both within the tumor and ex vivo by flow cytometry. T cell trafficking was blocked to examine changes to intratumoral T cells present at the time of vaccination.

**Results:** Vaccination led to increased intratumoral T cell numbers and delayed tumor growth. Expansion of T cells and tumor control did not require trafficking of T cells from the periphery. The increase in intratumoral T cells was associated with an acute burst in proliferation but not changes in viability. Intratumoral T cells had lower PD-1 and Eomes expression but were less functional after vaccination on a per cell basis. However, the increased intratumoral T cell numbers yielded increased effector T cells per tumor.

**Conclusions:** Pre-infiltrated CD8 T cells are responsive to CD40/TLR-mediated vaccination and sufficient for vaccination to delay tumor growth when additional T cell trafficking is blocked. This indicates that the existing T cell response and intratumoral DC could be critical for vaccination efficacy. This also suggests that circulating T cells may not be an appropriate biomarker for vaccination efficacy.

## BACKGROUND

Recent advances in checkpoint blockade and adoptive cell therapy demonstrate the potential for using adaptive immunity to fight cancer. Immune infiltration of tumors, especially by cytotoxic CD8 T cells, correlates with improved patient prognosis overall and in response to anti-PD1 therapy.(1,2) Therefore, boosting the presence of CD8 T cells in tumors is a major goal of cancer immunotherapy. One approach to induce CD8 T cell expansion in cancer patients is through vaccination. Early human vaccination trials were generally unsuccessful at controlling tumor growth, despite expanding antigen specific T cells.(3–7) More recent trials expanding the range of adjuvants or combining vaccination with other immunotherapies have improved the number of patients with stable or regressing tumors, but the magnitude of induced T cell expansion remains far below what is seen in response to infection in humans or with vaccination in tumor bearing mice.(8–11) As human vaccine trials evolve to incorporate next generation adjuvants and neoantigens, preclinical studies are needed to investigate the best combination of adjuvants for cancer vaccines and characterize their synergistic effects on immune responses.

In vivo, interactions with mature antigen presenting cells, primarily dendritic cells (DC), lead to CD8 T cell activation. One of the key drivers of DC maturation and subsequent T cell activation is stimulation of CD40 on DC, through engagement with CD40L on CD4 T cells.(12) While toll like receptor (TLR) agonists show some ability to promote T cell responses to vaccination,(13,14) the combination of CD40 stimulation and TLR agonists synergize to drive robust antigen specific CD8 T cell expansion.(15) The combination of αCD40 and TLR agonists has been shown to drive CD8 T cell expansion through upregulation of CD70 on DC(16,17) and to be dependent on IFNα/β.(11,15) Supporting the notion that activating DC promotes anti-tumor immunity, several studies demonstrate that vaccine approaches combining αCD40 agonistic antibody with TLR agonists are effective in mouse models of cancer.(11,18–21)

Despite the efficacy of vaccination with αCD40, polyinosinic-polycytidylic acid (polyI:C) and antigen in murine tumor models, the mechanistic basis for tumor control is still unclear. Some studies suggest that CD40-mediated activation of myeloid cells is sufficient for tumor control,(22–24) while others demonstrate that αCD40 vaccination increases antigen specific CD8 T cell numbers in the spleen and blood, and tumor control is dependent on CD8 T cells.(11,25) The contribution of CD4 T cells to tumor control after vaccination with αCD40, polyI:C, and antigens containing MHC class II epitopes is relatively unexplored. Whether vaccination with αCD40 and polyI:C alters the number and function of intratumoral T cells is also unknown. Interestingly, while studies show that IFNγ production by various effector cells substantially contributes to tumor control in multiple model systems, tumor control with αCD40, polyI:C, and peptide antigen vaccination is dependent on perforin but independent of IFNγ, suggesting that increased cytotoxicity may drive tumor control.(11) The subsets of immune cells with improved anti-tumor cytotoxicity after vaccination with αCD40 and polyI:C have not been well characterized. Further, the relative contribution of pre-existing tumor infiltrating T cells as compared to newly primed T cells arriving from the periphery to vaccination induced tumor control is unclear.

Here, using a well-established murine model of melanoma, we have examined T cell responses within the tumor after vaccination with αCD40, polyI:C, and tumor antigen. We show that either CD8 or CD4 T cells are sufficient for vaccination to delay tumor growth. We observe dramatically more CD8 and CD4 T cells in vaccinated tumors when trafficking is allowed and remarkably also when trafficking is blocked, indicating that vaccination promotes local expansion of T cells. Although vaccination expands T cells in the peripheral lymphoid tissues, we demonstrate that vaccination slows tumor growth without trafficking of additional T cells from the periphery. Both CD8 and CD4 T cells are more proliferative acutely after vaccination, although both T cell subsets do not maintain this increased proliferation and are ultimately not better able to survive within the tumor. Surprisingly, CD8 T cells were acutely less functional in situ after vaccination, although CD4 T cells were more functional after ex vivo stimulation. This reduction in function occurred even when new T cells were allowed to flux into tumors.However, the increased number of T cells counteracted their diminished function, ultimately leading to equivalent or higher numbers of functional T cells in the tumors of vaccinated mice. Therefore, vaccination limits tumor growth by sustaining the already infiltrated CD8 T cells within the tumor, without requiring additional T cell infiltration.

## METHODS

### Tumor model

The B16cOVA tumor line was previously generated in our laboratory.(26) B16cOVA cells (4×10^5^) were injected subcutaneously into the shoulder of C57BL/6 mice (National Cancer Institute). Tumors were measured by calipers in two directions and reported as tumor area. Mice were euthanized to harvest tumors at indicated time points or upon reaching the maximum allowed size. All mice were treated in accordance with policies established by the University of Virginia Animal Care and Use Committee.

### Treatments

Tumor bearing mice were vaccinated i.p. with 100 µg αCD40 (FGK45; BioXcell), 75 µg polyI:C (Invivogen), and 500 µg ovalbumin (Sigma) or 200 µg OVA_257-264_ (Genscript) at day ten after tumor injection. FTY720 (Sigma) was provided in the drinking water (2 µg/mL) starting at day nine after tumor implantation and supplemented with daily i.p. injection (25 µg) on days nine through twelve. Depletion antibodies (250 µg) for CD8 (2.43; BioXCell) and/or CD4 (GK1.5; BioXCell) were administered i.p. at day eight after tumor injection and again every four days for the remainder of the experiment. FTY720 and depletion efficiency were confirmed by flow cytometric analysis of blood in each experiment. LFA blocking antibody (M17/4; BioXCell) was administered i.p. (200 µg) every two days. Brefeldin A (Thermo Fisher) was injected i.v. (250 µg) five hours prior to harvest to block cytokine secretion. For OT1 transfer, spleens were harvested from OT1 mice originally from Taconic and maintained at the University of Virginia.

### Sample preparation

Tumors were excised and subjected to manual homogenization. T cells were isolated by density gradient separation with Lympholyte-m (Cedarlane). For in vitro function, isolated T cells were stimulated with plate-bound αCD3 (1µg/mL; eBioscience) or OVA_257_ peptide pulsed, CD45-mismatched splenocytes. H-2K^b^/SIINFEKL dextramer (Immudex) was used to identify OVA_257_ specific CD8. Antibodies and reagents used for staining are listed in table S1.

### Data analysis

Flow samples were collected on Cytoflex S (Beckman Coulter) or Attune NxT (Thermo Fisher) cytometers and analyzed using FlowJo V10. Statistical analyses were performed using Graphpad Prism V8. Tumor outgrowth data is presented as mean±SEM from one representative experiment and analyzed by Holm-Sidak multiple t tests. All other data is presented as mean±SD and analyzed by Welch’s t test for two groups or Holm-Sidak multiple t tests for more than two groups or time points. In some cases, pooled data is normalized to the control samples within each experiment to compensate for batch effects and shown as the relative expression of indicated factors. Differences were considered significant when p<0.05.

## RESULTS

### Vaccination is effective in established murine melanoma

To assess the mechanism by which vaccination with a CD40 agonistic antibody, TLR3 agonist polyI:C, and tumor antigen controls tumor growth, we utilized the B16cOVA melanoma model, which allowed us to assess the immune response to the well-defined pseudo-neoantigen ovalbumin (OVA) expressed by the tumor. When implanted subcutaneously, B16cOVA elicits a modest inflammatory immune response, including both CD8 and CD4 T cells, that diminishes over time and ultimately fails to control the tumor.(27) This provides the opportunity to improve both the function and number of infiltrating T cells. When tumor bearing mice were vaccinated three and ten days after tumor implantation by i.p. injection of either whole OVA protein or the CD8 immunodominant peptide OVA_257-264_ (SIINFEKL) in combination with agonistic αCD40 and polyI:C, mice were able to substantially control tumor growth (figure S1). To ask whether this immunization regimen was effective in established tumors and if tumor control required booster immunizations, we gave a single vaccination of αCD40, polyI:C, and ovalbumin once tumors were established at ten days post implantation. This treatment approach resulted in significantly smaller tumors compared to controls (figure 1A), affirming the efficacy of the vaccination approach. However, a single vaccination at day ten was insufficient to abolish tumor growth, and therefore provided the opportunity to investigate how vaccination alters the antitumor immune response.

**Figure 1.**
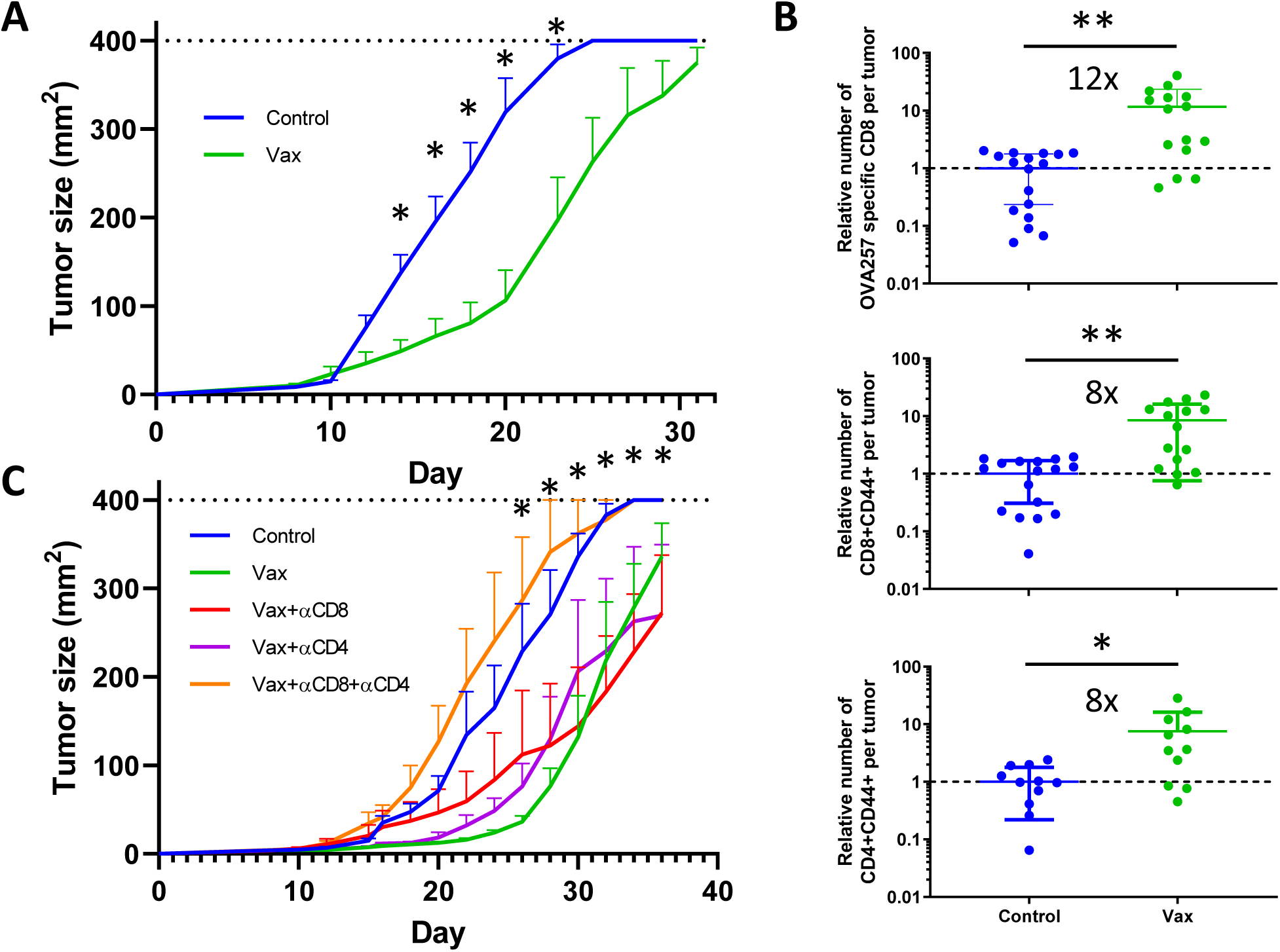
Therapeutic efficacy of vaccination with αCD40/polyI:C/antigen in murine melanoma. (A) Tumor size in mice bearing B16cOVA tumors vaccinated i.p. with αCD40/polyI:C/ovalbumin on day 10. n=6 mice per group. (B) Relative number of OVA_257_ specific CD8 T cells identified by MHC I dextramer (top) and total antigen experienced CD8 (middle) and CD4 (bottom) T cells infiltrating tumors at day 21 from mice vaccinated as in (A). Data are pooled from five experiments and normalized to the number of T cells in unvaccinated control tumors within each experiment. (C) Tumor size in mice vaccinated at day 10 as in (A) and injected i.p. with depleting antibodies to CD8 and/or CD4 every four days starting at day 8. n=4-7 mice per group. Data were analyzed by Holm-Sidak multiple t tests (AC) or Welch’s t test (B). (C) Significant difference between Control/Vax, Control/Vax+αCD8, and Control/Vax+αCD4 at indicated time points. *p<0.05, **p<0.01.

The antigen component of the vaccine includes epitopes recognized by both CD8 and CD4 T cells, so we measured the number of both intratumoral T cell subsets. Initial studies were performed at eleven days post-vaccination (day 21 after tumor implantation), when we observed reduced tumor size and sufficient time has passed for T cells to be activated and traffic to the tumors. Vaccination increased the number of OVA_257_ specific CD8 (12 fold) as well as total antigen experienced CD8 (8 fold) and CD4 (8 fold) T cells present in the tumors (figure 1B). Since vaccination with the OVA protein increased both CD8 and CD4 intratumoral T cells, we used depleting antibodies to test the relative contribution of CD8 versus CD4 T cells to delaying tumor growth. Tumor control after vaccination was lost when both CD8 and CD4 T cells were depleted, but either subset alone was sufficient to delay tumor growth (figure 1C). Thus, both CD8 and CD4 T cells can contribute to the control of tumor after αCD40 and polyI:C mediated vaccination with antigen containing epitopes for both subsets.

### Additional T cell infiltration is not required for vaccination to slow tumor growth

As the spontaneous anti-tumor immune response in untreated mice has a limited ability to control B16cOVA outgrowth, we would predict that vaccination slows tumor growth through the recruitment of more functional T cells to tumors. Alternatively, vaccination could lead to expansion and/or improved function in the T cells that have already infiltrated the tumor, despite their apparent exhausted phenotype in this model.(28) To test the contribution of new T cell infiltration to tumor control, we used the sphingosine-1-phosphate receptor agonist FTY720 to block T cell egress from lymphoid tissues and thereby block new T cell trafficking into tumors (figure 2A,S2).(29) Delaying FTY720 treatment allowed the initial antitumor T cell response to infiltrate the tumor prior to vaccination. Contrary to our expectation, vaccination still delayed tumor growth when FTY720 was used to block trafficking of additional T cells into the tumor (figure 2B). Separately, we prevented new T cell trafficking into tumors with αLFA1, which blocks T cell extravasation through LFA1 interaction with ICAM on vasculature.(30) Tumors in mice treated with αLFA1 and vaccinated grew slower than those treated with αLFA1 alone (figure S3A). Consistent with reports that αLFA1 alters T cell expansion and function in addition to blocking trafficking,(31,32) we observed decreased expansion of antigen specific T cells in the spleens of vaccinated tumor bearing mice (figure S3B). Despite the diminished T cell expansion, αLFA1 did not prevent vaccination from delaying tumor growth. Together, these data show that this vaccine can delay tumor growth through T cells that infiltrated the tumor prior to treatment without the need for recruitment of de novo activated T cells.

**Figure 2.**
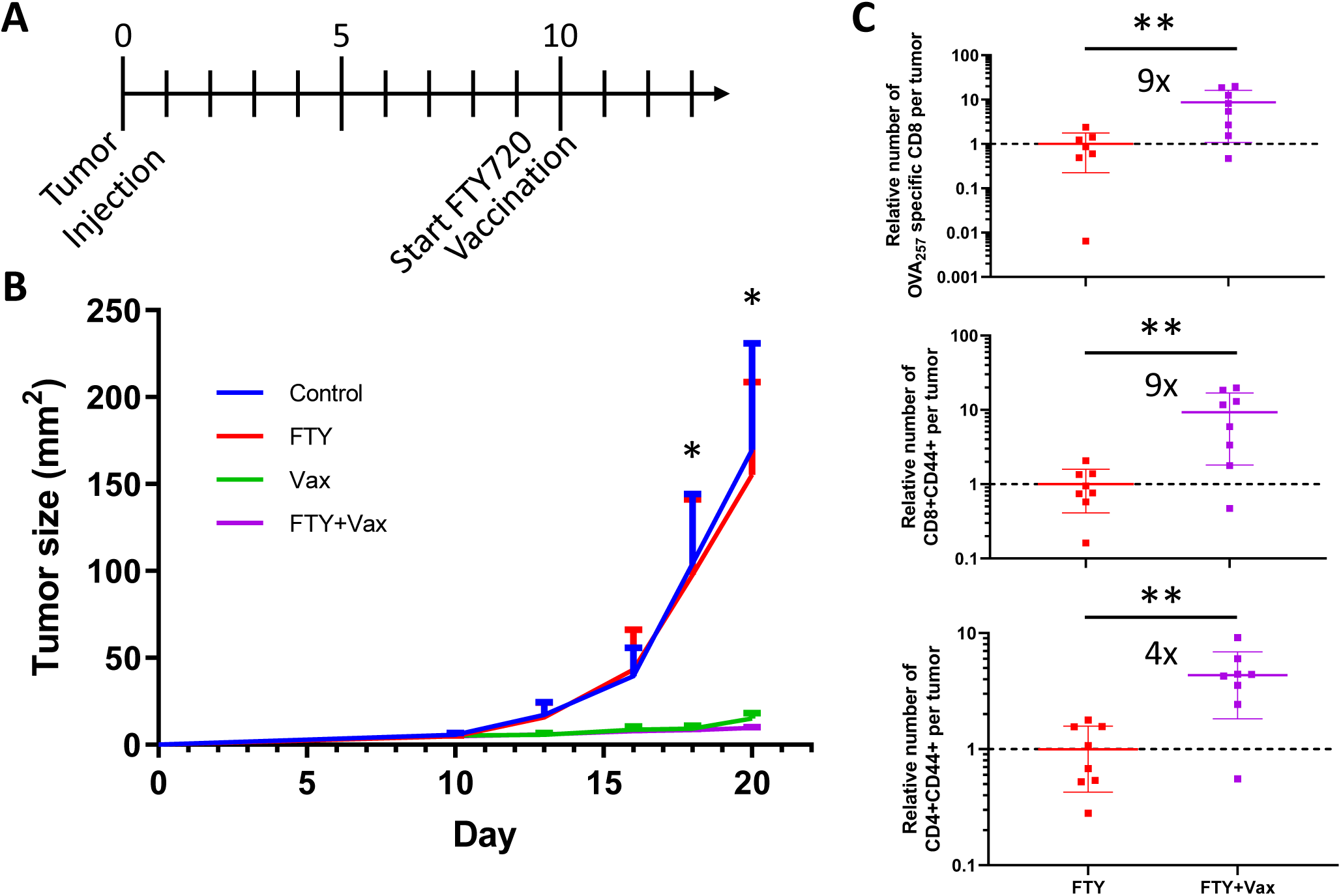
Additional T cell infiltration is not required for vaccination to slow tumor growth. (A) Mice bearing B16cOVA tumors were vaccinated and treated with FTY720 as indicated in the timeline and described in the methods. (B) Tumor size of mice treated with vaccination and FTY720. n=5 per group. (C) Relative number of OVA_257_ specific CD8 T cells (top) and total antigen experienced CD8 (middle) and CD4 (bottom) T cells infiltrating tumors at day 21. Data are pooled from 2 experiments. Each data point is from an individual mouse. Data were analyzed by Holm-Sidak multiple t tests (B) or Welch’s t test (C). (B) Significant difference between Control/Vax, Control/Vax+FTY, FTY/Vax, and FTY/FTY+Vax at indicated time points. *p<0.05, **p<0.01.

The vaccines ability to slow tumor growth through the pre-infiltrating T cells suggests that vaccination is acting within the tumor. In support of this, we found that intratumoral DC expressed significantly more CD86 two days after vaccination and only expressed IL12 (p40) in vaccinated mice (figure S4). Thus, intratumoral DC may serve as the intermediary linking vaccination and tumor control dependent on the already infiltrating T cells, and is consistent with previous reports demonstrating that intratumoral DC are critical contributors to vaccination-induced tumor control.(33,34)

### Vaccination sustains T cells within the tumor

We next considered how vaccination could slow tumor growth without new T cell infiltration. The two most likely explanations are that vaccination expanded the T cells within the tumor, or that vaccination improved the function of the T cells within the tumor. The number of OVA_257_ specific (9-fold) and total antigen experienced (9-fold) intratumoral CD8 T cells was higher at day eleven after vaccination when trafficking was limited with FTY720 (figure 2C). The number of CD4 T cells was also increased in vaccinated tumors (4-fold), although to a lesser extent than the CD8 T cells (figure 2C). This indicates that vaccination expanded or preserved both CD8 and CD4 T cells within the tumor. Surprisingly, we found similar numbers of all three groups of intratumoral T cells after vaccination with or without blocking T cell trafficking (figure S5A). This finding, when considered with the subdued expansion of vaccine-specific T cells in the spleens of tumor-bearing mice (figure S6), indicates that vaccination primarily affects T cells within the tumor rather than driving additional T cell infiltration.

To further understand how vaccination increased the number of both CD8 and CD4 T cells without new infiltration, we measured the number of intratumoral T cells from two to eleven days after vaccination while trafficking was blocked with FTY720. Whereas untreated tumors had fewer OVA_257_ specific CD8 T cells as time progressed, the number of OVA_257_ specific CD8 T cells remained steady in the vaccinated tumors (figure 3A). Although there were fewer CD4 T cells than OVA_257_ specific CD8 T cells, they followed the same pattern: sustained numbers in vaccinated tumors and a decline over time in unvaccinated tumors (figure 3E). To determine how vaccination sustained intratumoral T cell numbers, we measured proliferation and survival of the T cells. There was an increase in the proportion of OVA specific CD8 expressing Ki67 at four days after vaccination but a decrease in Ki67 expression at seven and eleven days after vaccination, indicating a transient increase in proliferation (figure 3B). Similarly, more CD4 T cells expressed Ki67 at four days, and fewer at eleven days after vaccination (figure 3F). To examine survival, we measured expression of Bcl2 and caspase 3/7 activation. Both OVA_257_ specific CD8 and total CD4 T cells had no difference in Bcl2 expression at two and four days after vaccination and a trending decrease in at seven days after vaccination (figure 3C,G). Similar activated caspase 3/7 levels indicated no difference in cell death in either OVA_257_ specific CD8 or total CD4 T cells at seven days after vaccination (figure 3D,H). Together, the Bcl2 and caspase 3/7 data indicate that vaccination does not improve intratumoral T cell survival. In summary, vaccination drives increased proliferation of intratumoral T cells acutely, which may contribute to sustained numbers of both CD8 and CD4 T cells within the tumor.

**Figure 3.**
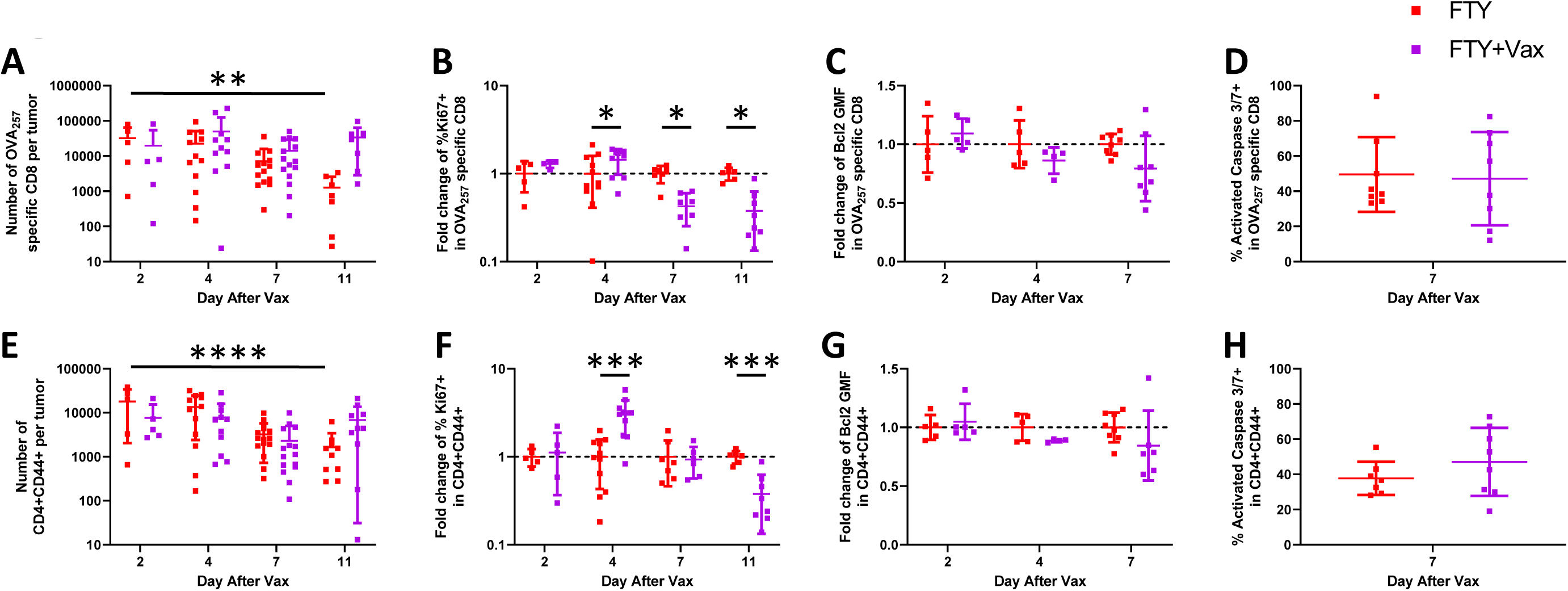
Vaccination sustains T cells within the tumor. Mice bearing B16cOVA tumors were vaccinated at day 10 and began receiving FTY720 at day 9 as described in methods and shown in figure 2A. Tumors were harvested and T cells were analyzed at 2, 4, 7, or 11 days after vaccination or from control mice at matching time points. (AE) Number of intratumoral OVA_257_ specific CD8 (A) and CD4 (E) T cells. (BF) Relative proportion of Ki67 expressing OVA_257_ specific CD8 (B) and CD4 (F) T cells within control or vaccinated tumors. (CG) Relative per cell expression of Bcl2 in intratumoral OVA_257_ specific CD8 (C) and CD4 (G) T cells. (DH) Relative caspase 3/7 activation in intratumoral OVA_257_ specific CD8 (D) and CD4 (H) T cells. Data were analyzed by ANOVA with test for linear trend (AE) or Holm-Sidak multiple t tests(B-D,F-H). *p<0.05, **p<0.01, ***p<0.001, ****p<0.0001.

### Vaccination does not improve T cell function within the tumor

Aside from T cell infiltration, we also predicted that vaccination would improve the function of intratumoral T cells. Supporting this, 11d after vaccination we found that the intensity of expression of the inhibitory receptor PD-1, which can lead to diminished T cell function when engaged by its ligands PD-L1 or PD-L2 on tumor or other immune cells in the tumor microenvironment, was significantly lower (∼2-fold) on a per cell basis on OVA_257_ specific CD8 T cells from vaccinated mice (figure S7A). To test function, we isolated OVA_257_ specific CD8 T cells from vaccinated and unvaccinated tumors and co-cultured them in vitro with OVA_257_-pulsed antigen presenting cells. Contrary to our expectations, significantly fewer OVA_257_ specific CD8 T cells from the tumors of vaccinated mice produced IFNγ and Granzyme B than those present in the tumors of control mice (figure S7B,C). Thus, diminished PD-1 expression did not lead to increased effector function of OVA_257_ specific CD8 T cells after vaccination. We also compared T cell function after vaccination when trafficking was uninhibited. Remarkably, fewer OVA_257_ specific CD8 T cells from vaccinated mice produced IFNγ than from unvaccinated mice when trafficking was allowed (figure S5B), although Granzyme B expression was not significantly different (figure S5C). This suggests that after vaccination newly infiltrating T cells are more cytotoxic but not more capable of IFNγ production compared to T cells already present within the tumor.

An inflection point in tumor growth occurs around ten days after vaccination, which could suggest that the effects of a single vaccination are only temporary. Therefore, we examined the function of both CD8 and CD4 T cells at earlier time points after vaccination. We used FTY720 to block trafficking of newly primed T cells and harvested tumors at 4 and 7 days after vaccination. In this instance, we gave brefeldin A (BFA) i.v. prior to harvest to block cytokine secretion in situ and thus measure cytokine production by T cells within the tumors. While there was no difference in effector activity in T cells present in the tumors of vaccinated and control mice at day four, a lower proportion of OVA_257_ specific CD8 produced IFNγ (∼50% reduction) and Granzyme B (∼30% reduction) within the tumor seven days after vaccination (figure 4A-C). As in vivo BFA reports steady state cytokine production and may reflect lack of encounter with antigen-bearing cells, we also measured the T cell functional activity by stimulating the T cells isolated from tumors with a low concentration of αCD3 ex vivo to directly engage the T cell receptors. There was a trend for a higher frequency of OVA_257_-specific CD8 T cells expressing IFNγ at day four but, as seen with in situ assessment, a significant reduction at day seven after vaccination (figure S8A). Ex vivo stimulation also allowed us to assess the capability of the T cells to degranulate, which is required for release of cytotoxic molecules such as Granzyme B. There was no difference in the capability of OVA_257_ specific CD8 T cells to degranulate as measured by surface exposure of CD107a during the ex vivo stimulation (figure S8B). This suggests that vaccination did not alter the cytotoxicity of OVA_257_ specific CD8 T cells. Additionally, we saw the same pattern of decreased effector function in intratumoral CD8 T cells that were not specific for OVA_257_ after vaccination, both in situ and ex vivo (not shown). This demonstrates that vaccination had similar effects on all intratumoral CD8 T cells regardless of the actual antigen they recognized, and further supports that vaccination is primarily affecting the intratumoral T cells. Together, this data shows that vaccination is not controlling tumor growth by improving CD8 T cell effector molecule expression or degranulation.

**Figure 4.**
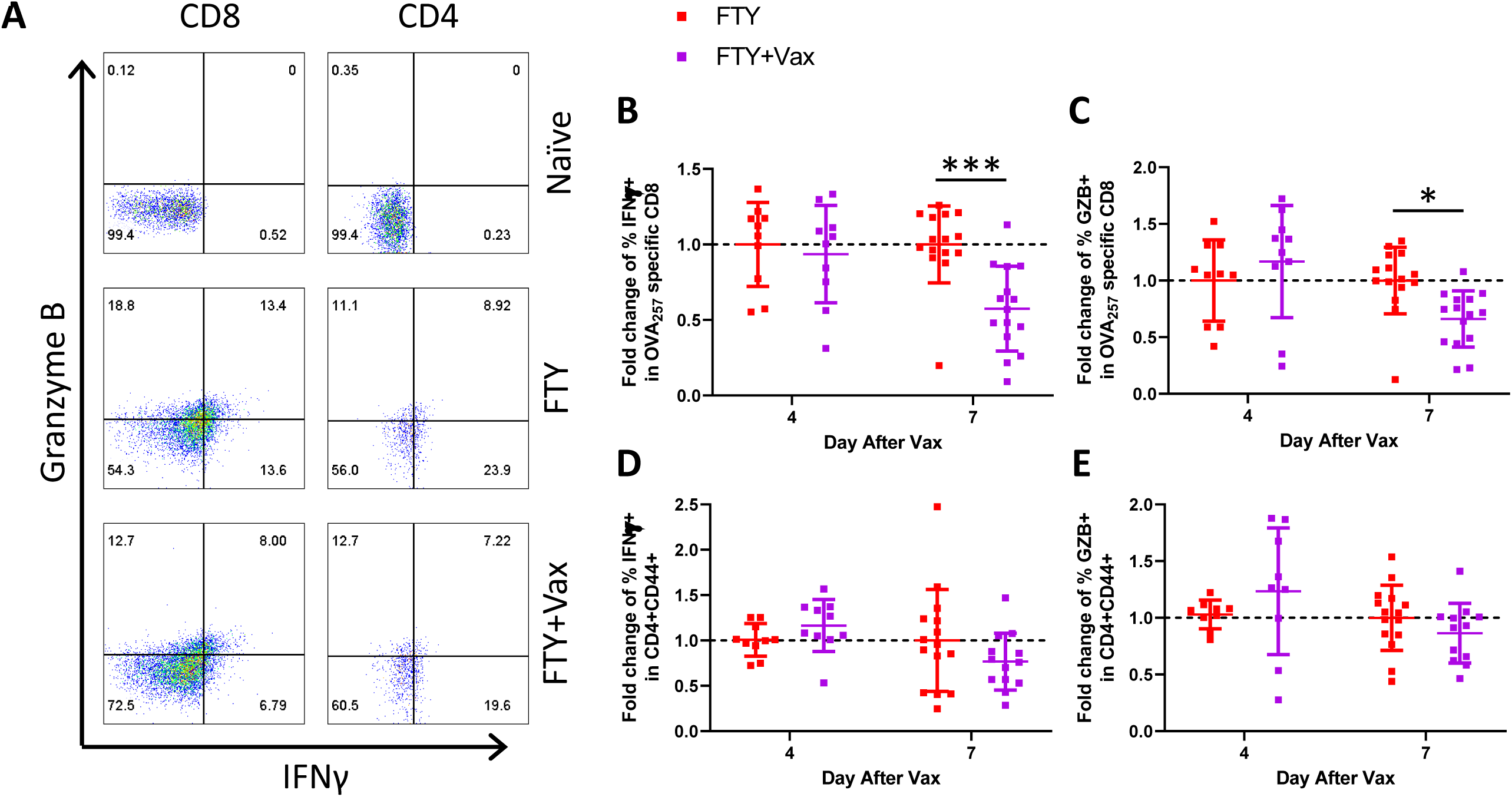
Vaccination does not improve T cell function within the tumor. Mice bearing B16cOVA tumors were vaccinated at day 10 and began receiving FTY720 at day 9 as described in methods and shown in figure 2A. Brefeldin A was administered 5 hours prior to harvest to measure in vivo expression of IFNγ and Granzyme B by tumor infiltrating OVA_257_ specific CD8 and CD4 T cells. (A) Representative flow plots of naïve T cells from the spleen (top) and OVA_257_ specific CD8 or CD4 T cells from control (middle) and vaccinated (bottom) tumors. (BC) Relative proportion of OVA_257_ specific CD8 T cells expressing IFNγ (B) and Granzyme B (C) from 3 pooled experiments. (DE) Relative proportion of CD4 T cells expressing IFNγ (D) and Granzyme B (E) from 3 pooled experiments. Data were analyzed by Holm-Sidak multiple t tests. *p<0.05, **p<0.01, ***p<0.001.

While CD8 T cells are important based on their cytotoxic capabilities, CD4 T cells can contribute to the antitumor response through cytokines, and in some cases killing of target cells. Therefore, we also examined the function of CD4 T cells within the tumor. In situ, intratumoral CD4 T cells had no significant difference in either IFNγ or Granzyme B expression at day four or day seven after vaccination compared to controls (figure 4D,E). Intratumoral CD4 T cells also showed no difference in degranulation based on surface exposure of CD107a during ex vivo stimulation with αCD3 (figure S8D). In contrast to both OVA_257_ specific CD8 and CD4 T cell in situ production of IFNγ, more CD4 T cells expressed IFNγ at both four and seven days after vaccination when stimulated ex vivo (figure S8C). Although the vaccinated CD4 T cells did not make more IFNγ within the tumor, they have increased potential to contribute effector cytokines.

Together these data show that vaccination differentially affects CD8 and CD4 T cell function, and that these effects are discrete when examined in the steady state as compared to after restimulation. Importantly, the relative increase in the number of CD8 and CD4 T cells within the tumor found after vaccination substantially outweighs the reduction in function observed in situ, resulting in equivalent or significantly more effector T cells within the tumors of vaccinated mice (figure S9A-F), especially at d11 post vaccination.

### Vaccination does not expand T cell subsets identified by Eomes or Tcf1 expression

We were intrigued by the reduced function of the vaccinated TIL. Recently, populations of exhausted CD8 T cells that can proliferate and generate more functional CD8 T cells have been separately identified based on two transcription factors: low expression of Eomes and high expression of Tcf1.(35–37) Therefore, we asked whether a subset of intratumoral CD8 T cells identified by Eomes or Tcf1 expression was responding to vaccination. We found decreased Eomes expression in OVA_257_ specific CD8 T cells from vaccinated tumors at day 17 (figure 5A). However, unlike published data, we found that T cells with higher Eomes expression had the highest degree of proliferation, and this was true in both control and vaccinated mice (figure 5B). Further, we found that T cells with higher Eomes expression had increased IFNγ and Granzyme B expression, irrespective of vaccination (figure 5C,D). Tcf1 expression declined in OVA_257_ specific CD8 T cells between day 14 and 17, but Tcf1 expression was not impacted by vaccination (figure 5E). As with Eomes, we found higher Ki67 expression in Tcf1+ cells compared to Tcf1-cells, as expected, and this relationship was also unaltered by vaccination (figure 5F). We also found that Tcf1 expression correlated with higher IFNγ and Granzyme B expression, in contrast to previous reports (figure 5G,H). In summary, higher expression levels of Eomes and Tcf1 expression correlate with more functional intratumoral CD8 T cells. However, the increase in intratumoral T cells that accompanies vaccination does not result from expansion of a subset of intratumoral CD8 T cells expressing Eomes or Tcf1.

**Figure 5.**
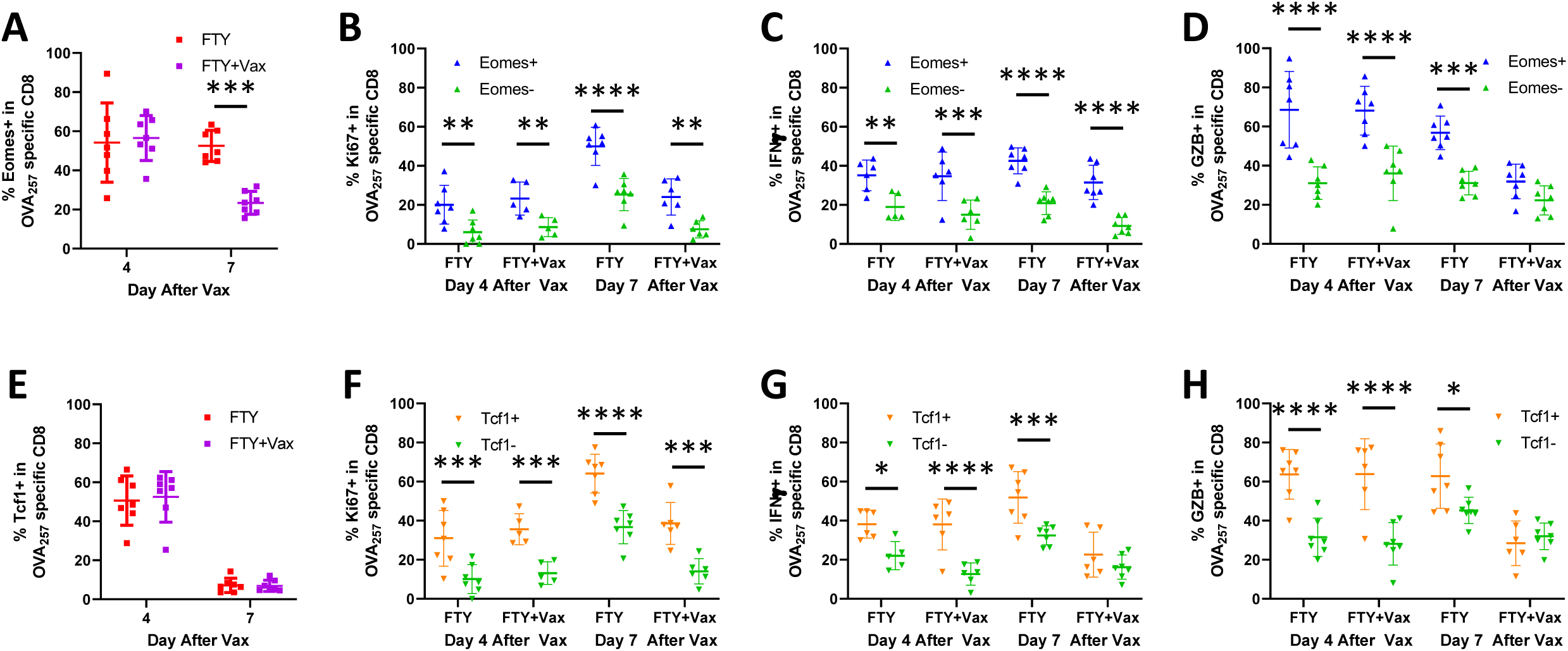
Vaccination does not improve proliferation or function in a CD8 T cell subset identified by Eomes or Tcf1. Mice bearing B16cOVA tumors were vaccinated at day 10 and began receiving FTY720 at day 9 as described in methods and shown in figure 2A. Brefeldin A was administered 5 hours prior to harvest. (A) Expression of Eomes in intratumoral OVA_257_ specific CD8 T cells. (B) Expression of Ki67 in in intratumoral OVA_257_ specific CD8 T cells that do (blue) or do not (green) express Eomes. (CD) Proportion expressing IFNγ (C) and Granzyme B (D) of intratumoral OVA_257_ specific CD8 T cells that do (blue) or do not (green) express Eomes. (E) Expression of Tcf1 in intratumoral OVA_257_ specific CD8 T cells. (F) Expression of Ki67 in in intratumoral OVA_257_ specific CD8 T cells that do (orange) or do not (green) express Tcf1. (GH) Proportion expressing IFNγ (G) and Granzyme B (H) of intratumoral OVA_257_ specific CD8 T cells that do (orange) or do not (green) express Tcf1. Data were analyzed by Holm-Sidak multiple t tests. *p<0.05, **p<0.01, ***p<0.001, ****p<0.0001.

### Pre-infiltrated CD8 T cells are sufficient for vaccination to delay tumor growth

Although vaccination sustained both CD8 and CD4 T cell numbers within the tumors, there were more OVA_257_ specific and total CD8 T cells per tumor (figure 3A,E). However, CD8 had decreased function in situ after vaccination whereas CD4 T cells were more functional ex vivo (figure 4,S8). Further, when T cell trafficking was unrestricted, either CD8 or CD4 T cells were sufficient to slow tumor growth (figure 1C). Since additional T cell infiltration was not required for vaccination to delay tumor growth (figure 2A), we next assessed if both the pre-infiltrated CD8 and CD4 T cells had the potential to impede tumor growth. To test this, we combined T cell depletion with FTY720 to block trafficking into the tumor (figure 6A). As expected, based on our earlier depletion experiment, depleting both CD8 and CD4 T cells in FTY720 treated mice resulted in the loss of vaccination induced tumor control (figure 6B). When we depleted only CD4 cells in FTY720 treated mice, vaccination still delayed tumor growth. However, when we blocked trafficking with FTY720 and depleted CD8 T cells, the already infiltrating CD4 T cells were insufficient to slow tumor growth after vaccination. Thus, when T cell trafficking to tumor is prevented, only the pre-infiltrated CD8 T cells could slow tumor growth. Although the intratumoral CD8 T cells exhibited lower levels of effector functions after vaccination, they still had an essential role. Thus, the CD8 T cells infiltrating the tumor at the time of vaccination play critical contributions to the ability of vaccination to delay tumor growth.

**Figure 6.**
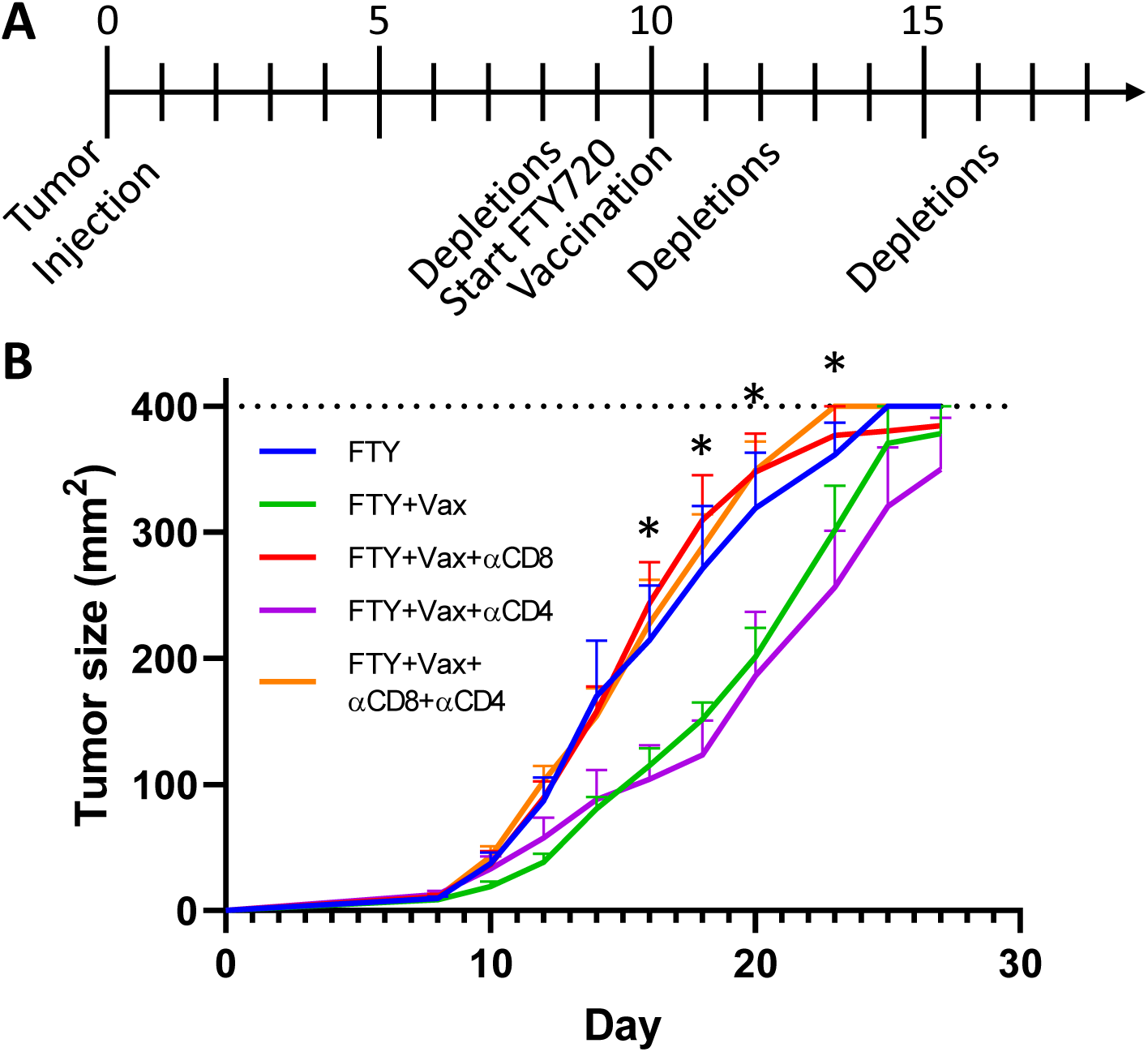
Pre-infiltrated CD8 T cells are sufficient for vaccination to delay tumor growth. (A) Mice bearing B16cOVA tumors were treated with vaccination, FTY720, and depleting antibodies as indicated in the timeline and described in the methods. (B) Tumor size of mice treated as described in (A). n=6-7 per group. Data were analyzed by Holm-Sidak multiple t tests with significant difference between FTY/FTY+Vax and FTY/FTY+Vax+αCD4 at indicated time points. *p<0.05.

## DISCUSSION

Previously, and in our data, vaccination regimens with αCD40, polyI:C, and antigen control tumor growth and generate large T cell responses observed in the spleen and blood. As T cells are known to become exhausted within the tumor, it was assumed that newly activated T cells were trafficking to the tumor and responsible for controlling tumor growth. However, the relative increase in the numbers of T cells in tumors after vaccination was greater than seen in the periphery, and this relative increase was sustained when trafficking from the periphery was blocked. Further, by blocking trafficking we have shown that the existing CD8 T cells are sufficient for vaccination to slow tumor growth without additional T cell infiltration. Together, these data make the unexpected argument that pre-existing intratumoral T cells are critical responders to this vaccine regimen.

Despite the compelling evidence that vaccination leads to tumor control via pre-existing intratumoral T cells, there are several paradoxes concerning the mechanistic basis of tumor control. First, we predicted that vaccination would lead to more functional T cells within the tumor. Instead, a lower proportion of CD8 T cells from vaccinated tumors expressed IFNγ and Granzyme B in situ acutely after vaccination, and they were no better at degranulating. While a greater proportion of CD4 T cells from vaccinated tumors had the potential to produce more IFNγ ex vivo, they were also not more functional in situ within the tumor. Intriguingly, ex vivo activity more closely correlated with tumor control, suggesting that relatively few tumor-specific T cells are encountering antigen at any given time within tumors or that they are being actively suppressed. The basis for the loss of effector activity by the CD8 tumor infiltrating lymphocytes 7-11d after vaccination remains unclear. OVA_257_ specific CD8 T cells expressed less PD-1, which we would expect to improve function within the tumor. Decreased Eomes expression may have contributed to diminished function. However, T-bet, which was highly expressed (not shown), and Tcf1 levels were unaffected by vaccination. Glycolytic activity in T cells has been shown to regulate IFNγ translation,(38) and we have previously shown that tumor infiltrating lymphoyctes have limited glycolysis.(28) We are currently investigating whether vaccination leads to a further reduction in glycolytic activity in intratumoral T cells. Notably, intratumoral T cells from vaccinated mice with intact trafficking were also less functional than from control mice, suggesting that loss of functional activity occurs within the tumor microenvironment. Critically, however, the decreased function of intratumoral T cells was offset by vaccination sustaining T cell numbers within the tumors, resulting in an equivalency or increase in the number of cytokine expressing T cells per tumor, even when gauged by in situ function (figure S9A,B,D,E). Thus, the impact of vaccination on T cell persistence in the tumor mitigates their diminished functionality. It is possible that other effector mechanisms or aspects of T cell biology that we currently do not fully understand are contributing to tumor control.

The second paradox concerns the capacity of vaccination to sustain the number of both CD8 and CD4 T cells present in the tumor even when trafficking of T cells into the tumor is blocked. This suggests, surprisingly, that the sustained intratumoral T cell numbers are a function of maintaining or expanding the existing T cells rather than new T cell trafficking to tumors. While there was increased proliferation of intratumoral T cells acutely after vaccination, T cells within the tumor were less proliferative at later time points. Nevertheless, there could be a subset of T cells proliferating in response to vaccination that is missed by considering the total population. To address this possibility, we focused on two transcription factors known to be critical for T cell proliferation in a tumor setting, Eomes and Tcf1. Expression of both Eomes and Tcf1 correlated with increased proliferation, yet vaccination did not affect Tcf1 and led to reduced Eomes expression in intratumoral CD8 T cells, and did not improve proliferation in a Tcf1+ or Eomes+ subset. T cells from vaccinated mice expressed less of the survival protein Bcl2, but had equivalent caspase 3/7 activation. Together, these data suggest neither expansion of less differentiated T cells nor increased survival were responsible for sustaining T cell numbers in response to vaccination. We also considered retention of T cells in the tumor. However, we found that the majority of OVA specific T cells expressed the retention integrin CD103 with no difference after vaccination (not shown). Thus, we currently ascribe the impact of vaccination on T cell numbers to the acute burst in proliferation, but why proliferation is rapidly curtailed remains to be determined.

These mechanistic paradoxes are important as αCD40 and TLR3 agonists are being aggressively developed for clinical implementation and suggest that efficacy may be divorced from traditional biomarkers of tumor control. A combination of agonistic αCD40, the polyI:C derivative polyIC:LC, and peptides generated a substantial antigen specific CD8 response in nonhuman primates.(39) CD40 monoclonal antibodies have shown to be safe in human trials, with limited efficacy on tumor control as a single agent,(40–42) and better response rates when combined with chemotherapy or checkpoint blockade.(43,44) PolyIC:LC has been incorporated into human vaccines that generate T cell responses but not necessarily control tumor growth(8,9). Two clinical trials with αCD40, polyIC:LC, and peptide antigens have been initiated for melanoma patients (NCT03597282, NCT04364230), but have not reported outcomes. Based on our data, the existing T cell response may be critical for such vaccination regimens to be effective. Additionally, our data suggests that circulating T cells are not a definitive biomarker for vaccination efficacy, as they are not required for tumor control.

In our model, vaccination was administered at a site distant to the tumor, yet CD8 T cells that were already present within the tumor were sufficient for tumor control. This suggests that vaccine administration directly into the tumor could have the same effects within the tumor, and likely at much lower doses, which could limit systemic toxicity. Additionally, since vaccination affected T cells within the tumor, the antigen component may not be necessary. However, when we treated mice with αCD40 and polyI:C without additional antigen plus FTY720 to block trafficking, only half the mice were able to delay tumor growth (not shown). Mice that responded had equivalent tumor control as mice that received the full vaccination, suggesting that the amount of tumor-derived antigen presented by intratumoral DC is insufficient in non-responder mice. This argues for including either shared antigens or neoantigens identified by sequencing tumors in human vaccination regimens. Alternatively, treatments that liberate tumor antigens could be combined with αCD40 and polyIC:LC, and this may account for the cases where αCD40 and chemotherapy were successfully combined in patients. Future studies in these areas would provide additional clarity and potential effectiveness for translating vaccination withαCD40 and polyI:C to human cancer patients.

## CONCLUSIONS

Vaccination with αCD40, polyI:C, and tumor antigen can delay the growth of established melanoma tumors by proficiently sustaining pre-infiltrating CD8 T cells. Intratumoral T cells lose effector function with time after vaccination, yet are more acutely proliferative resulting in a relative increase in the number of effector T cells within the tumors of vaccinated mice. Although vaccination drives T cell expansion in the periphery, our data shows that additional T cell infiltration is unnecessary for tumor control, and that the existing intratumoral CD8 T cells therefore critically contribute to tumor control after this vaccination regime.

## ACKNOWLEDGEMENTS

The authors would like to thank Dr. Alexandra Witter and Ms. Marissa Gonzales for assistance and discussions and the Beirne B. Carter Center for Immunology Research for use of the flow cytometry instruments.

---

**Figure S1**. Therapeutic efficacy of double vaccination with αCD40/poly(I:C)/antigen in murine melanoma. (A) Tumor size in mice bearing B16cOVA tumors vaccinated i.p. with αCD40/polyI:C/ovalbumin on day 3 and 10 or no vaccination control. n=4-5 mice per group. (A) Tumor size in mice bearing B16cOVA tumors vaccinated i.p. with αCD40/polyI:C/OVA_257-264_ on day 3 and 10 or no vaccination control. n=3-4 mice per group. (AB) Data were analyzed by Holm-Sidak multiple t tests. *p<0.05.

**Figure S2**. FTY720 blocks T cell trafficking through blood. B16cOVA tumor bearing mice were treated with FTY starting at day 9 and/or vaccinated with αCD40/polyI:C/ovalbumin on day 10. Number of CD8 (A) and CD4 (B) T cells per 100 ul of blood at day 13. Data were analyzed by Welch’s t test. *p<0.05.

**Figure S3**. Effects of αLFA1 on tumor growth and CD8 T cell expansion. (A) Tumor size in mice bearing B16cOVA tumors vaccinated i.p. with αCD40/polyI:C/ovalbumin on day 10 and/or αLFA1 every two days starting at day 9. n=6-8 mice per group (B) Approximately 10,000 OT1 T cells were transferred into mice bearing B16cOVA tumors on day 9. Mice were treated with αLFA1 on day 9, 11, 13, and 15 and/or immunized with αCD40/polyI:C/ovalbumin on day 10. Spleens were harvested and OT1 T cells were counted at day 17. Data were analyzed by Holm-Sidak multiple t tests (A) or Welch’s t test (B). (A) Significant difference between Control and Vax+LFA at indicated time points. *p<0.05.

**Figure S4**. Vaccination leads to acute intratumoral DC maturation. B16cOVA tumor bearing mice were treated with FTY starting at day 9 and/or vaccinated with αCD40/polyI:C/ovalbumin on day 10. Brefeldin A was administered 5 hours prior to harvest to measure cytokine expression within the tumor. Intratumoral DC were identified as CD11c+MHCII^hi^ cells. (AB) Expression of CD86 by proportion (A) and per cell expression (B) on intratumoral DC. (C) In vivo expression of IL12 by intratumoral DC. (A-C) Data are representative of two experiments. Data were analyzed by Welch’s t test. *p<0.05, **p<0.01.

**Figure S5**. Number and function of intratumoral T cells with and without blocked trafficking. B16cOVA tumor bearing mice were treated with FTY starting at day 9 and/or vaccinated with αCD40/polyI:C/ovalbumin on day 10. (A) Relative number of OVA_257_ specific CD8 T cells (left) and total antigen experienced CD8 (middle) and CD4 (right) T cells infiltrating tumors at day 21. (BC) Expression of IFNγ (B) and Granzyme B (C) by tumor infiltrating OVA_257_ specific CD8 T cells after in vitro stimulation with OVA_257_ pulsed splenocytes. (A-B) Data are pooled from two experiments. (A-C) Data were analyzed by Welch’s t test. *p<0.05.

**Figure S6**. Vaccination expands antigen specific T cells in naïve and tumor bearing mice. Mice were vaccinated with αCD40/polyI:C/ovalbumin in naïve mice or ten days after B16cOVA tumor implantation. Number of OVA_257_ specific CD8 T cells present in the spleen seven days after vaccination. Non-tumor and tumor bearing mice were vaccinated in separate experiments. All groups were significantly different from each other based on Welch’s t test with Dunnet’s correction.

**Figure S7**. Function of OVA_257_ specific CD8 T cells at day 11 after vaccination. Mice bearing B16cOVA tumors were vaccinated at day 10 and began receiving FTY720 at day 9 as described in methods and shown in figure 2A. (A) Geometric mean fluorescence of PD1 on tumor infiltrating OVA_257_ specific CD8 T cells. (BC) Proportion of tumor infiltrating OVA_257_ specific CD8 T cells expressing IFNγ (B) and Granzyme B (C) after in vitro stimulation with OVA_257_ pulsed splenocytes. (A-C) Data are pooled from 2 experiments. Each data point is from an individual mouse. Data were analyzed by Welch’s t test. *p<0.05, **p<0.01.

**Figure S8**. Ex vivo function of intratumoral T cells. Mice bearing B16cOVA tumors were vaccinated at day 10 and began receiving FTY720 at day 9 as described in methods. Intratumoral T cells were isolated and stimulated with αCD3 for four hours. (AB) Relative proportion of OVA_257_ specific CD8 T cells expressing IFNγ (A) and CD107a (B). (CD) Relative proportion of CD4 T cells expressing IFNγ (C) and CD107a (D). (A-D) Data are pooled from three experiments and normalized to the proportion of positive cells in unvaccinated control tumors within each experiment. Data were analyzed by Holm-Sidak multiple t tests. *p<0.05, **p<0.01.

**Figure S9**. Number of functional intratumoral T cells. Mice bearing B16cOVA tumors were vaccinated at day 10 and began receiving FTY720 at day 9 as described in methods and shown in figure 2A. Brefeldin A was administered 5 hours prior to harvest to measure in vivo expression of IFNγ and Granzyme B by tumor infiltrating OVA_257_ specific CD8 T cells (ABDE) or intratumoral T cells were isolated and stimulated in vitro for four hours (CF). (AB) Relative number of OVA_257_ specific CD8 T cells expressing IFNγ (A) and Granzyme B (B) in vivo. (C) Relative number of OVA_257_ specific CD8 T cells expressing IFNγ ex vivo. (DE) Relative number of CD4 T cells expressing IFNγ (D) and Granzyme B (E) in vivo. (F) Relative number of CD4T cells expressing IFNγ ex vivo. (A-F) Data were analyzed by Holm-Sidak multiple t tests. *p<0.05, **p<0.01, ***p<0.001, ****p<0.0001.

