## Supplementary figures and images for "Pre-existing intratumoral CD8 T cells substantially contribute to control tumors following therapeutic anti-CD40 and polyI:C based vaccination"

### Supplemental Figures

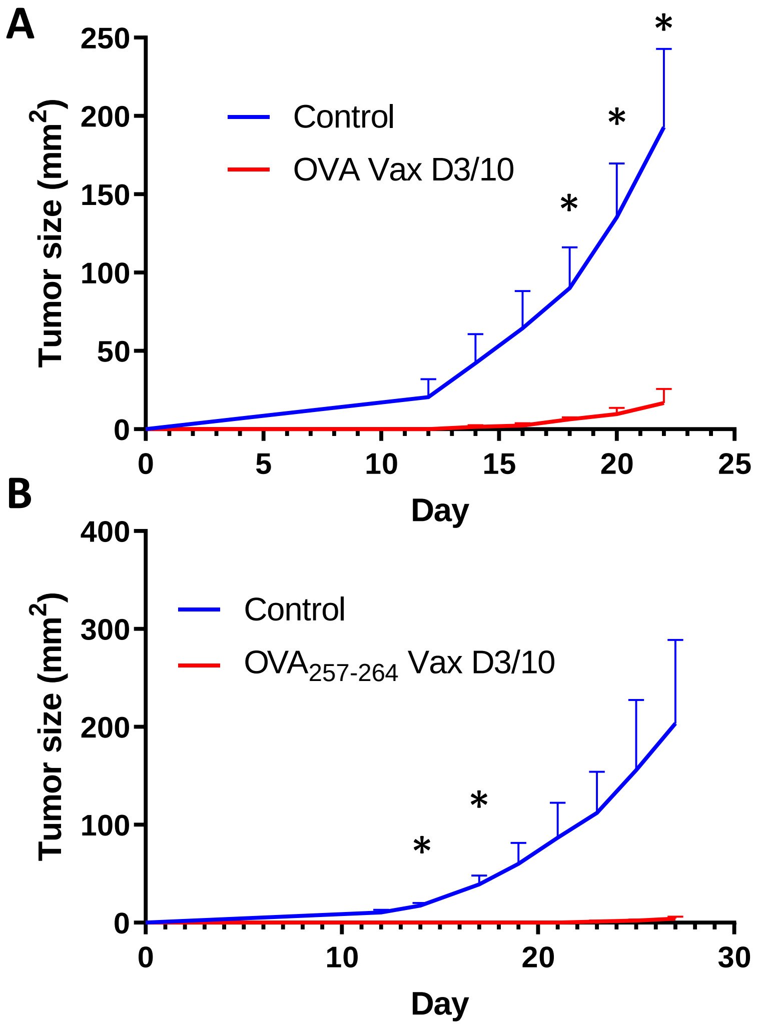

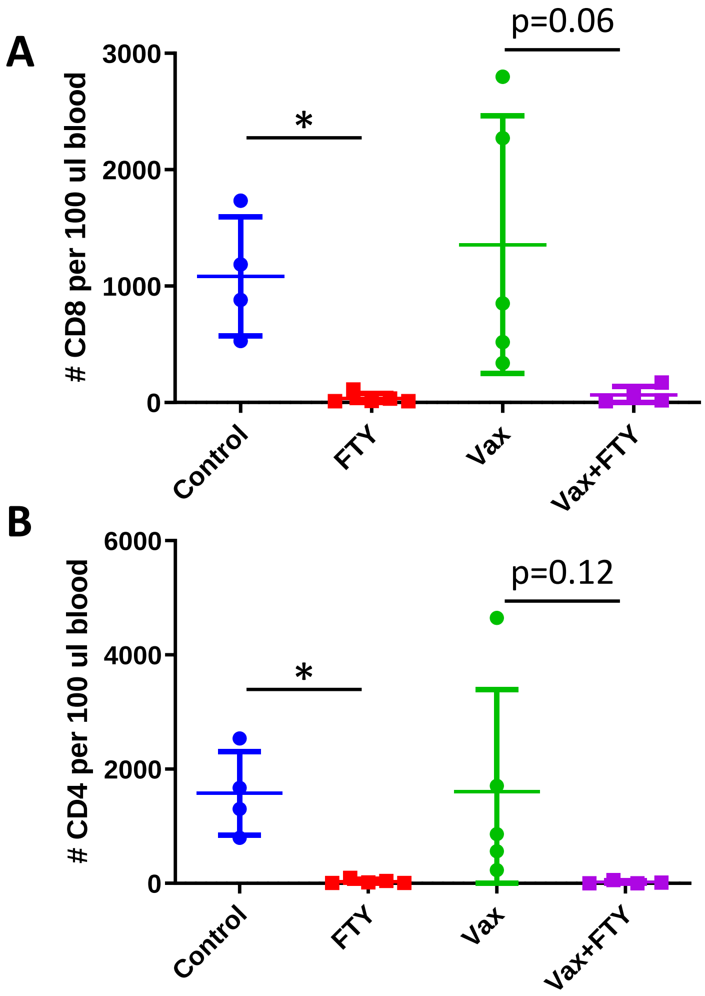

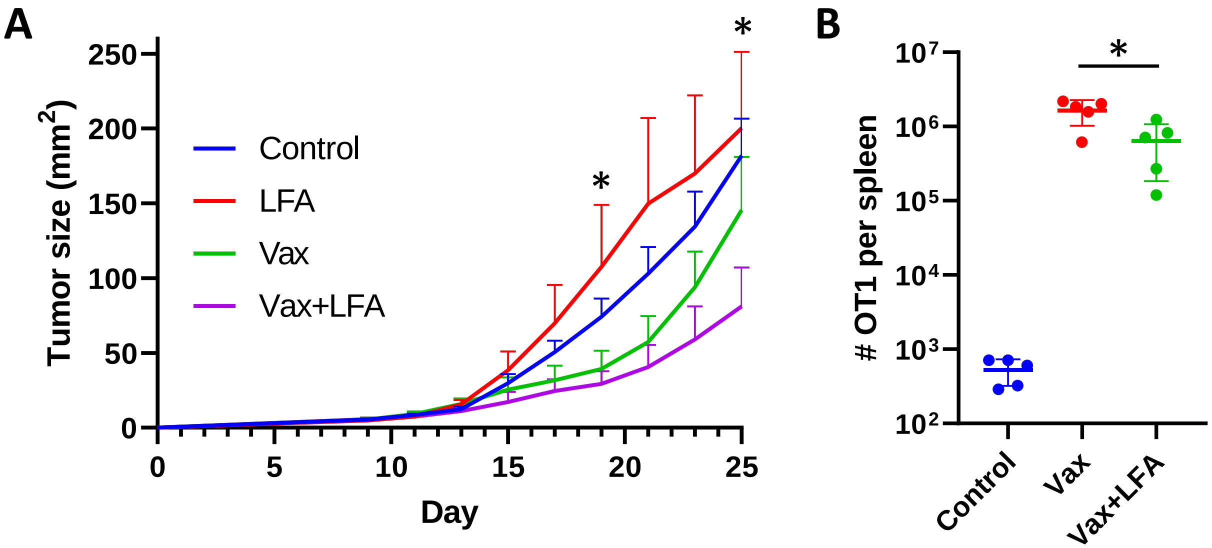

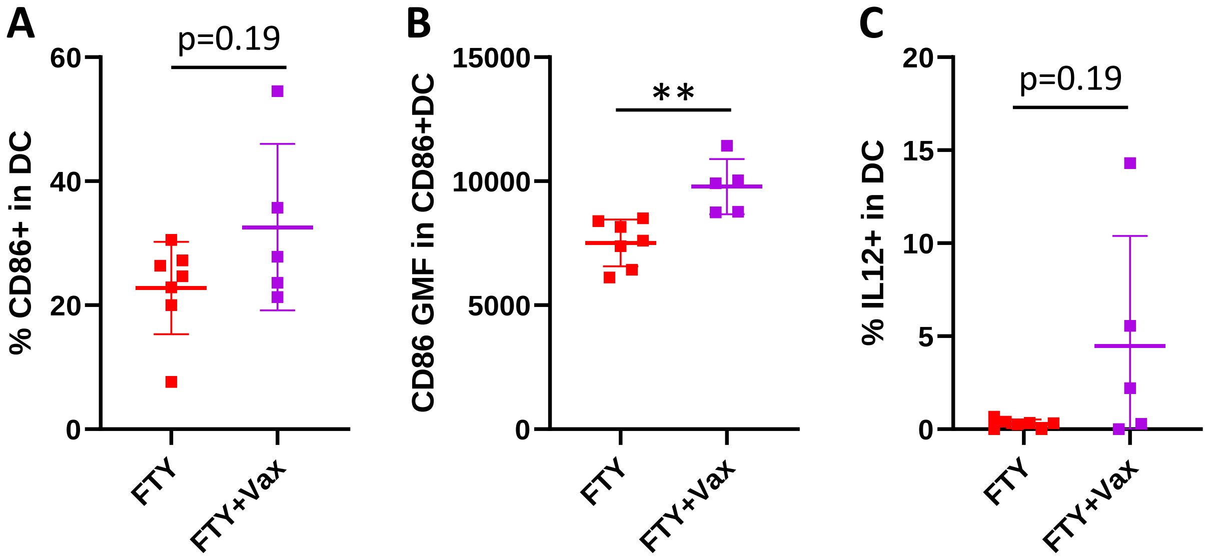

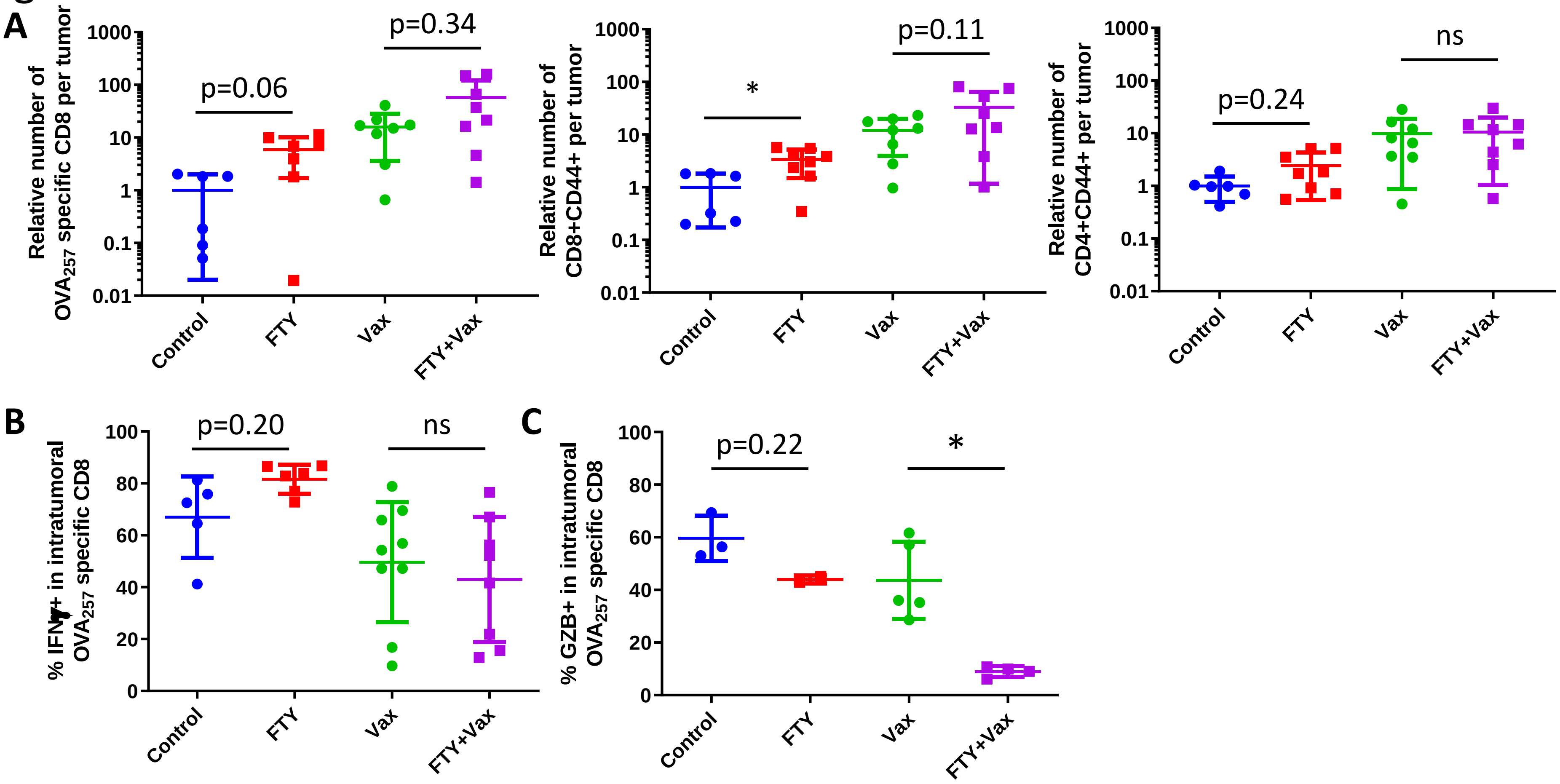

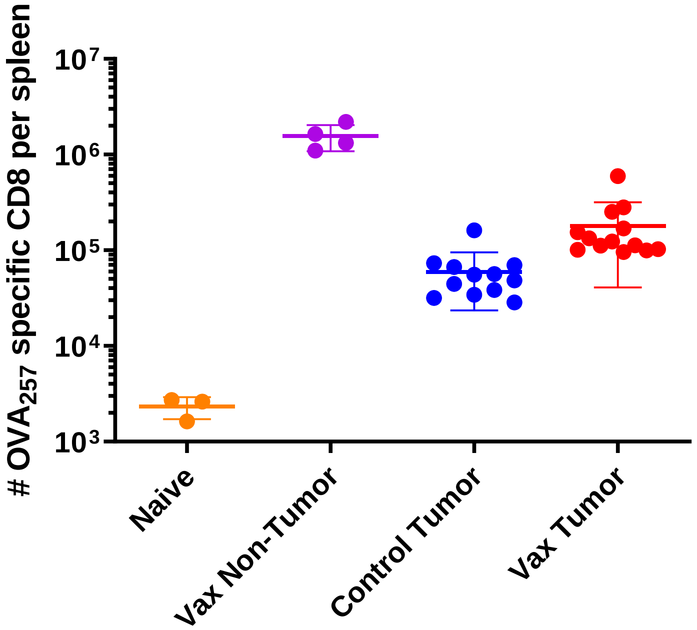

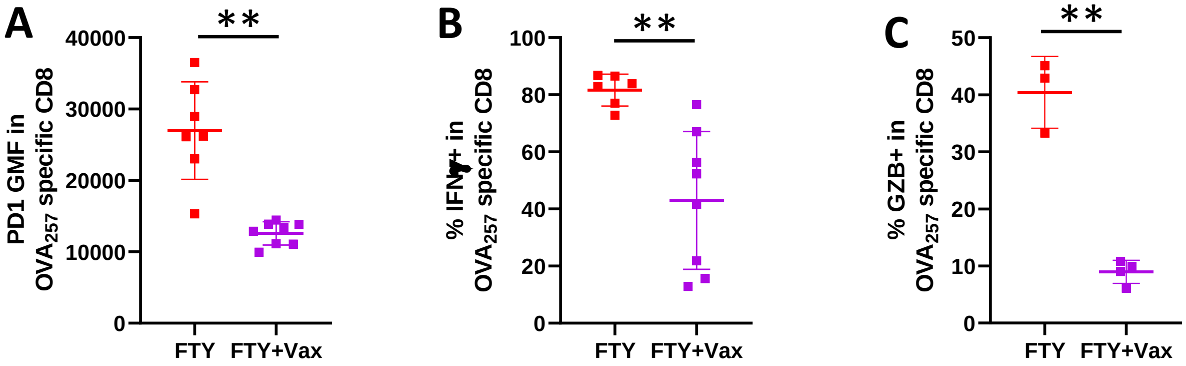

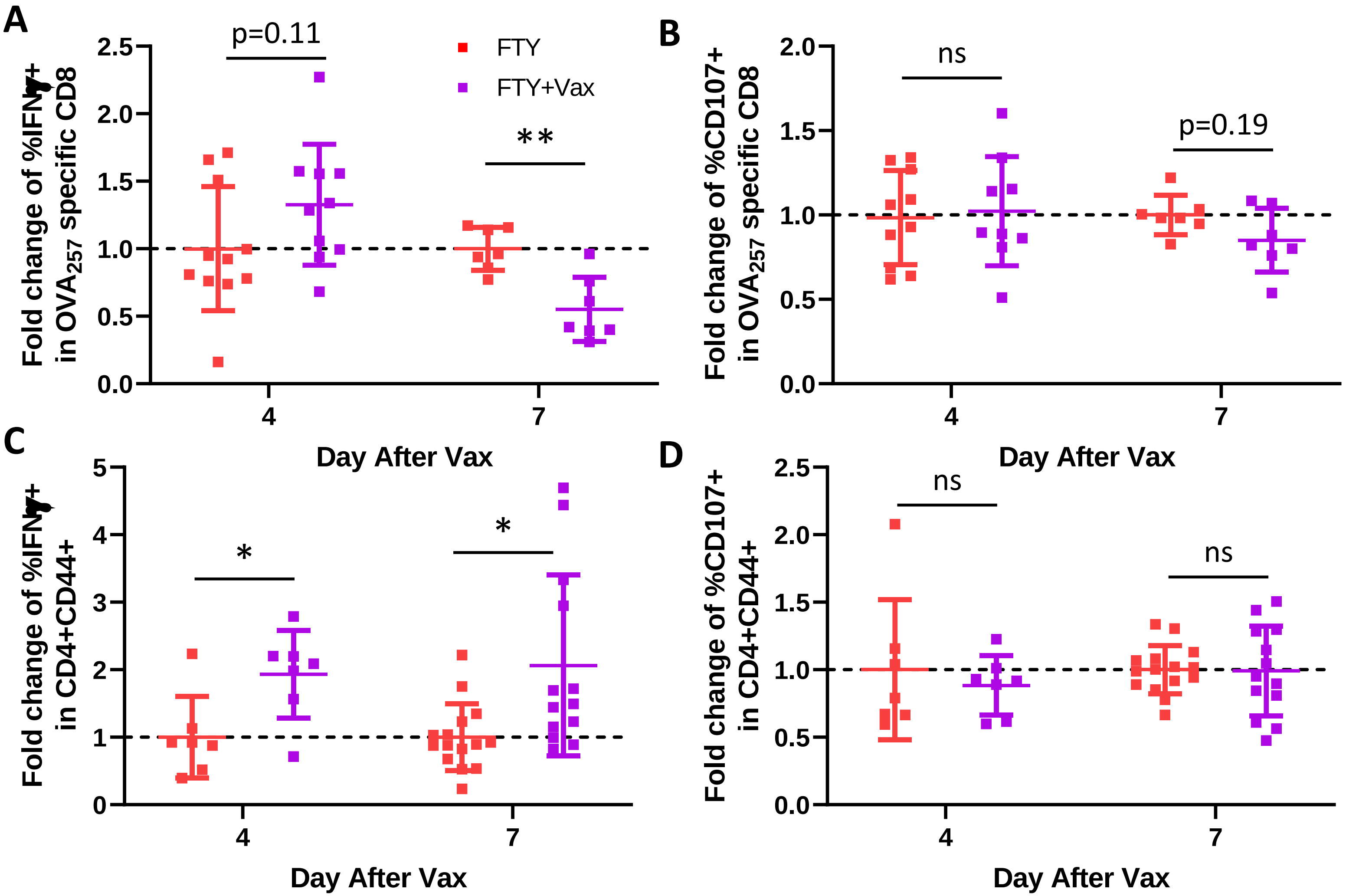

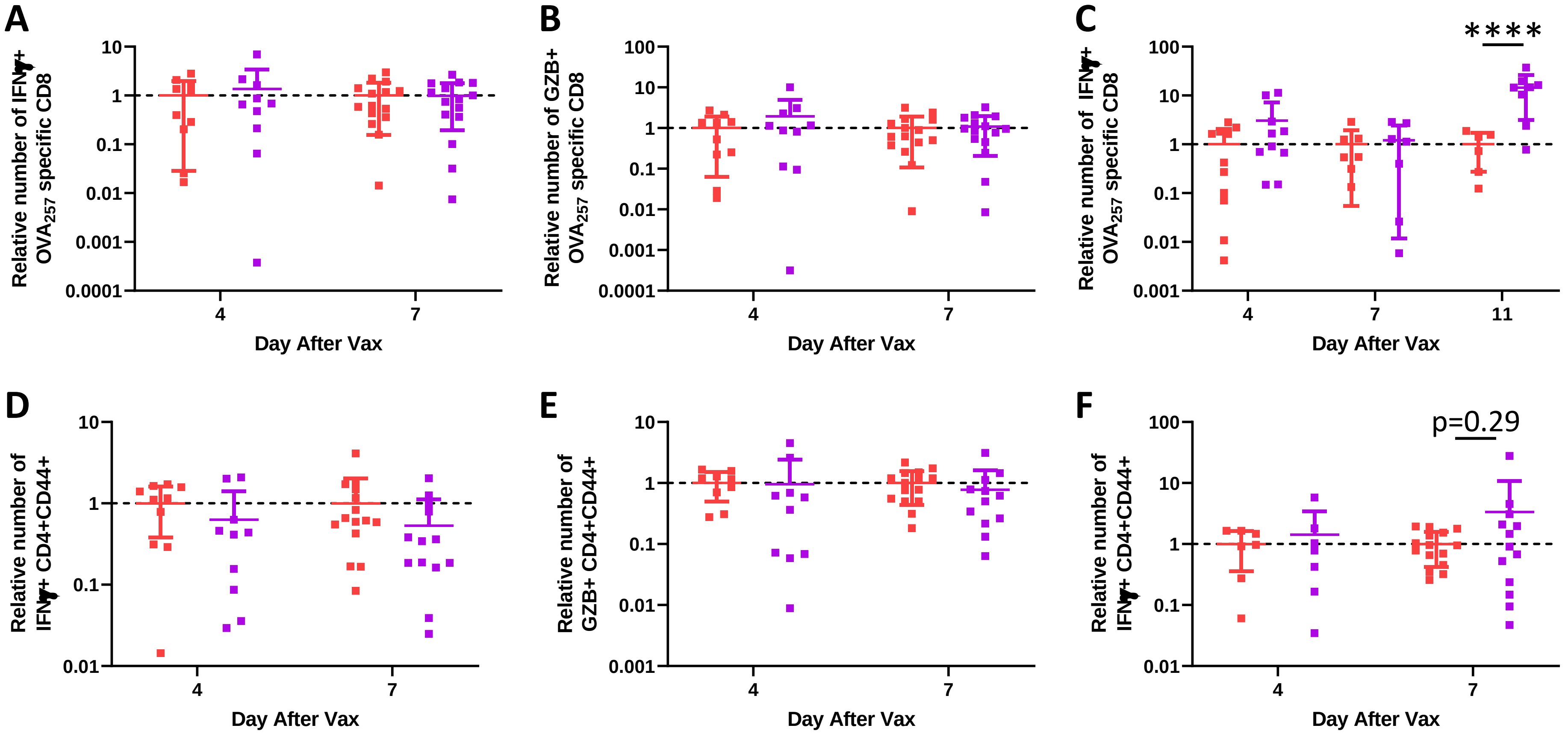
