## Supplementary material for "Pre-existing intratumoral CD8 T cells substantially contribute to control tumors following therapeutic anti-CD40 and polyI:C based vaccination": Materials Table

| Antibody | Fluorochrome(s) | Clone | Company |
|---|---|---|---|
| Live/Dead | Aqua | N/A | Life Technologies |
| Bcl2 | e450 | 10C4 | eBioscience |
| Caspase 3/7 | Green Detection Reage | N/A | Life Technologies |
| CD107a | PE-Cy7 | 1D4B | BD Biosciences |
| CD11c | APC, PE | N418 | eBioscience |
| CD4 | PE-594 | RM4-5 | BioLegend |
| CD44 | AF700 | IM7 | eBioscience |
| CD45.2 | APC-780, e450, FITC | 104 | eBioscience |
| CD8 | APC, BV650 | 53-6.7 | BioLegend |
| CD86 | BV650 | GL-1 | BioLegend |
| Eomes | PerCP-e710 | Dan11mag | eBioscience |
| Granzyme B | PE-Cy7 | NGZB | eBioscience |
| IFNγ | APC, PE-Cy7 | XMG1.2 | eBioscience |
| IL12 | PE | C15.6 | BD Biosciences |
| Ki67 | FITC, PE-Cy7 | SolA15 | eBioscience |
| MHC II | PE-Cy7 | M5/114.15.2 | eBioscience |
| OVA dextramer | PE | N/A | Immudex |
| PD1 | BV605 | 29F.1A12 | BioLegend |
| Tcf1 | AF488 | C63D9 | Cell Signaling |

| Population | Gating strategy | | | | | | |
|---|---|---|---|---|---|---|---|
| CD8 | Scatter | Singlet | Live | CD45.2+ | CD8+ | CD44+ | dextramer+ (when $OVA_{257}$ specific) |
| CD4 | Scatter | Singlet | Live | CD45.2+ | CD4+ | CD44+ | |
| DC | Scatter | Singlet | Live | CD45.2+ | CD11c hi | MHC II+ | |
